# Transcriptomic Analysis of the Newborn Mammary Gland Identifies Prenatal Ethanol-Induced Remodeling of Insulin Signaling and Enhanced Proliferative Programming

**DOI:** 10.64898/2026.09.22.753596

**Authors:** Zhikun Ma, Yongxuan Liu, Amanda B Parris, Xiaohe Yang

**Affiliations:** Julius L. Chambers Biomedical/Biotechnology Research Institute; Department of Biological and Biomedical Sciences, North Carolina Central University, North Carolina Research Campus, 500 Laureate Way, Kannapolis, NC, 28081, USA

**Author notes:** Correspondence: Xiaohe Yang Ph.D.

**Keywords:** prenatal alcohol exposure, mammary gland, transcriptomics, insulin signaling, IGF signaling, IRS1, developmental programming

## Abstract

Prenatal alcohol exposure (PAE) can produce persistent alterations in mammary development and increase susceptibility to mammary tumorigenesis, but the early molecular events underlying these effects remain poorly understood. Here, we examined whether PAE alters the molecular program of the mammary gland at birth. Pregnant MMTV-ErbB2 mice received control or ethanol-containing liquid diets during gestation, and mammary tissues were collected from newborn female offspring. RNA sequencing revealed exposure-level-dependent transcriptional reprogramming, with substantially broader alterations following moderate compared with lower ethanol exposure. Pathway analyses identified prominent changes in metabolic and growth-regulatory networks, including insulin signaling, with coordinated alterations in components of the insulin/IGF-IRS-PI3K/AKT pathway. Biochemical analyses further demonstrated exposure-dependent changes in IGFBP expression and IRS1 abundance and phosphorylation, supporting remodeling of insulin/IGF-associated signaling. Transcriptomic analyses also revealed enrichment of RNA metabolic and translational processes and, particularly following moderate exposure, enhanced DNA-replication, cell-cycle, and mitotic programs. These molecular signatures were accompanied by increased proliferative features in newborn mammary tissue. Together, these findings demonstrate that PAE establishes substantial molecular reprogramming of the mammary gland by birth and identify altered insulin/metabolic signaling and proliferative programming as prominent features of this early developmental response.

## Introduction

Breast cancer susceptibility is influenced not only by genetic and adult-life exposures but also by developmental events that establish the cellular and molecular properties of the mammary gland [1–5]. Mammary development begins during embryogenesis, when epithelial specification, mammary bud formation, and interactions with the surrounding mesenchyme establish the foundation for subsequent postnatal growth [6–8]. Perturbations during this sensitive developmental period may therefore alter mammary developmental trajectories and influence tissue function later in life, consistent with the broader concept of developmental origins of adult disease [9, 10].

Alcohol is a well-established developmental toxicant [11, 12]. This is exemplified by the development of Fetal Alcohol Spectrum Disorders (FASDs) [11, 13]. However, its effects on mammary development remain much less understood than those on the nervous system and other developing organs [13, 14]. Experimental studies have shown that prenatal alcohol exposure (PAE) can alter mammary development and increase susceptibility to mammary tumorigenesis later in life [15–17]. In rodent models, gestational ethanol exposure has been associated with changes in mammary growth, epithelial proliferation, hormonal and growth-factor signaling, and subsequent tumor development [16, 18–20]. These observations suggest that PAE may establish an altered developmental state within the mammary gland before later hormonal, environmental, or oncogenic challenges occur.

Our previous studies using the MMTV-ErbB2 mammary tumor model provided evidence supporting this possibility [15]. PAE altered mammary morphogenesis and estrogen receptor and growth-factor signaling and was associated with increased mammary tumor susceptibility [15]. More recently, studies in a non-tumorigenic mouse background demonstrated persistent effects of PAE on postnatal mammary morphogenesis, epithelial proliferation, mammary epithelial populations, stem/progenitor-associated activities, and growth-regulatory signaling [21]. Together with findings from other experimental models, these observations support the concept that a transient prenatal exposure can produce durable reprogramming of mammary development.

However, the molecular events established within the mammary gland during the prenatal exposure period itself remain poorly defined. Most previous studies have examined mammary tissue during postnatal development or evaluated tumor outcomes later in life, when the initial effects of PAE may have become integrated with subsequent endocrine and developmental influences. Mammary bud development begins during mid-gestation, and this period of rapid development and establishment of mammary epithelial/stem-progenitor populations may be particularly sensitive to environmental perturbations [22–24]. Based on previous studies from our group and others demonstrating persistent postnatal effects of PAE on mammary development, a critical question is whether these later phenotypes originate from molecular reprogramming established during fetal mammary development. Specifically, how extensively does the developing mammary gland respond to PAE at the transcriptional level, which biological programs are affected, and are molecular features associated with later developmental phenotypes already evident at birth? Analysis of the newborn mammary gland provides an opportunity to examine this early molecular state immediately following prenatal exposure and before extensive postnatal mammary morphogenesis.

In the present study, we extended our previous work [15], by using RNA-seq to examine gene-expression profiles in mammary tissues from newborn female MMTV-ErbB2 offspring and address the questions outlined above. Newborn mammary tissues from control and two PAE groups, lower exposure (PAE-L) and moderate exposure (PAE-M), exhibited substantial differences in gene expression, with bioinformatic analyses revealing exposure-level-dependent transcriptional reprogramming. Among the altered pathways, insulin/metabolic signaling was prominently affected, particularly in the PAE-M group, and was accompanied by exposure-dependent changes in IRS1 abundance and phosphorylation. PAE also altered transcriptional programs associated with RNA metabolism and translation and, particularly at the moderate exposure level, enhanced programs related to DNA replication, mitosis, and cell-cycle progression. These molecular changes were accompanied by increased proliferative features in newborn mammary tissue, as indicated by Ki67 staining [25]. Together, our findings demonstrate that PAE establishes molecular reprogramming of the mammary gland that is already evident at birth, characterized by altered insulin/metabolic signaling and enhanced proliferative programming, providing an early molecular context for the persistent mammary phenotypes associated with prenatal ethanol exposure.

## Materials and Methods

### Animals and prenatal ethanol exposure

FVB/N-Tg(MMTV-neu) mice were obtained from The Jackson Laboratory (Bar Harbor, ME, USA) and maintained in the institutional animal facility under a 12-h light/12-h dark cycle. Mice were provided an estrogen-free AIN-93G diet (Bio-Serv, Flemington, NJ) and water ad libitum except during the defined liquid-diet exposure period. All animal procedures were conducted under protocols approved by the Institutional Animal Care and Use Committee. Female and male mice were paired for breeding at approximately 8 weeks of age, and females were monitored for pregnancy. On gestational day (GD) 10, pregnant dams were randomly assigned to control, lower ethanol exposure (PAE-L), or moderate ethanol exposure (PAE-M) groups (10 dams/group). Beginning on GD11 and continuing through the remainder of pregnancy (GD11-20), PAE-L and PAE-M dams received Lieber–DeCarli ’82 liquid diets containing 1.7% or 3.4% (v/v) ethanol, respectively, corresponding to one-third or two-thirds of the 5% ethanol formulation. Lieber–DeCarli Shake and Pour formulated liquid diets were obtained from Bio-Serv (Flemington, NJ, USA). Control dams received the corresponding isocaloric ethanol-free liquid diet. Newborn female offspring were collected at birth, and mammary tissues were harvested immediately for subsequent analyses.

### Collection of newborn mammary tissue

Female offspring were euthanized at birth, and inguinal mammary glands were dissected under a stereomicroscope with minimal surrounding tissue. Samples were either processed immediately or snap-frozen in liquid nitrogen and stored at −80°C for molecular analyses, while tissues designated for imaging were prepared for cryosectioning. To minimize litter effects, biological replicates were obtained from different litters whenever possible, and offspring from multiple independent litters were distributed across experimental analyses to avoid disproportionate representation of any single litter.

### RNA isolation and RNA sequencing

Total RNA was isolated from frozen newborn mammary tissue using TRIzol reagent according to the manufacturer’s protocol. RNA concentration and purity were assessed spectrophotometrically, and RNA integrity was evaluated before sequencing. Three independent biological samples from each group (control, PAE-L, and PAE-M; *n* = 3/group) were used for RNA-seq analysis. RNA samples were submitted to Novogene for library preparation, sequencing, and initial bioinformatic processing. Briefly, polyadenylated RNA was enriched from total RNA, fragmented, converted to cDNA, and used to construct sequencing libraries. Libraries were quality controlled and sequenced on an Illumina platform. The resulting dataset contained approximately 40– 47 million raw reads per sample, with Q20 values of approximately 97.5–97.8%.

### RNA-seq processing and differential expression analysis

Raw sequencing reads were subjected to quality filtering to remove adapter-contaminated and low-quality reads. Clean reads were aligned to the mouse reference genome (mm10) using HISAT2. Across the nine samples, approximately 92–94% of clean reads mapped to the reference genome. Gene-level read counts were generated for expression analysis, and expression profiles were additionally represented as FPKM where appropriate for visualization and sample-level comparisons. Differential gene-expression analysis was performed using DESeq2 for the PAE-L versus control, PAE-M versus control comparisons. DESeq2 modeling was based on normalized read counts and a negative-binomial framework, with multiple-testing correction using the Benjamini-Hochberg procedure. For global descriptions and volcano plots, genes were evaluated according to the statistical criteria specified for each analysis. Adjusted *P* values were used for focused analyses requiring control of the false-discovery rate.

### Functional enrichment and pathway analysis

Functional interpretation of the RNA-seq data was performed using Gene Ontology (GO), Kyoto Encyclopedia of Genes and Genomes (KEGG), and Reactome pathway enrichment analyses. Initial enrichment analyses were performed using the clusterProfiler framework. Pathways with adjusted *P* < 0.05 were considered significantly enriched unless otherwise indicated. Because insulin-related pathways emerged prominently from the enrichment analyses, genes contributing to the KEGG insulin signaling pathway (mmu04910) were examined in greater detail. For the focused PAE-M versus control heatmap, pathway-associated genes meeting adjusted *P* < 0.05 and log2 fold change| ≥ 0.5 were selected. Expression values were standardized within each gene (row Z-score), followed by hierarchical clustering of genes; samples were displayed according to experimental group.

Reactome enrichment analysis was used independently to examine broader biological programs altered by prenatal ethanol exposure, with particular attention to cell-cycle, DNA-replication, RNA-processing, and translational pathways.

### Quantitative real-time PCR

Total RNA isolated from newborn mammary tissue was reverse transcribed to cDNA using iScript cDNA Synthesis Kit (Bio-Rad, CA, USA). Quantitative real-time PCR (qRT-PCR) was performed using All-in-One qPCR Mix (GeneCopoeia) and gene-specific primers. IGFBP3: F 5′-GTTGGGAGGGGAGGTAGGT-3′, R 5′-GAGCAGTACCCGCTGAGG-3′, IGFBP5: F 5′-GGAAGACCTTGGGGGAGTAG-3′, R 5′-TCAACGAAAAGAGCTACGGC-3′. The relative fold change of mRNA expression was determined for each group using the 2^-ΔΔCt^ method after normalization to the GAPDH.

### Western blot analysis

Newborn mammary tissues were homogenized on ice in protein lysis buffer containing protease and phosphatase inhibitor cocktails (Thermo scientific, IL). Protein concentrations were determined using the bicinchoninic acid (BCA) assay, and equal amounts of protein (30 μg/sample) were resolved by 10–12% SDS-PAGE. Proteins were transferred to nitrocellulose membranes, which were blocked for 1 h in Tris-buffered saline containing 0.1% Tween-20 and 5% nonfat dry milk. Membranes were incubated with primary antibodies overnight at 4°C, washed, and subsequently incubated with the appropriate HRP-conjugated secondary antibodies for 1 h. The target proteins were visualized using enhanced chemiluminescence reagents and captured using an Azure imaging system (Dublin, CA). Band intensity was quantified using ImageJ. Where phosphorylated proteins were examined, phosphoprotein abundance was normalized to the corresponding total protein. Total protein abundance was normalized to the indicated β-Actin loading control. Proteins examined included IRS1(#3407), phospho-IRS1 (Ser612) (#3203), phospho-IRS1 (Tyr895) (#3070), phospho-Akt1 (Ser473) (#4060), ErbB3 (#12708), phospho-ERK1/2 (Thr202/Tyr204) (#9101) were purchased from Cell signaling Technology, EGFR (sc-03), IGF1R (sc-712), Akt1 (sc-5298), Erk2 (sc-1647), ERα (sc-8002), Sox2 (sc-398254), IGFBP5 (sc-515116) and β-Actin (sc-47778) were purchased from Santa Cruz Biotechnology.

### Immunofluorescence analysis of newborn mammary tissue

Newborn mammary tissues were embedded in optimal cutting temperature compound and cryosectioned at 5 μm. Sections were mounted on positively charged slides, equilibrated to room temperature, washed with PBS, and permeabilized with 0.1% Triton X-100. Nonspecific binding was blocked for 1 h with 5% normal serum containing 1% bovine serum albumin. Sections were incubated overnight at 4°C with antibodies against Ki67 (1:1,000) and α-smooth muscle actin (α-SMA; 1:300). After washing, appropriate fluorophore-conjugated secondary antibodies were applied for 1 h at room temperature in the dark. Nuclei were counterstained with DAPI, and slides were mounted in antifade medium. Fluorescence images were acquired using a Nikon fluorescence microscope.

For quantitative analysis of proliferation, Ki67-positive nuclei were quantified within the mammary epithelial structures identified by α-SMA staining and expressed relative to the total number of DAPI-positive nuclei.

### Statistical analysis

Data from biochemical, qRT-PCR, and imaging experiments are presented as mean ± SEM unless otherwise indicated. Individual offspring were treated as biological replicates, with litter representation controlled as described above. Comparisons involving three experimental groups were analyzed using one-way ANOVA followed by an appropriate multiple-comparison test; two-group comparisons were analyzed using an unpaired two-tailed Student’s *t*-test where applicable. Statistical significance was defined as *P* < 0.05. RNA-seq statistical procedures and multiple-testing correction were performed as described above.

## Results

### 1. Prenatal ethanol exposure induces broad transcriptomic alterations in the newborn mammary gland

To determine whether prenatal alcohol exposure (PAE) alters the molecular program of the developing mammary gland, pregnant mice were maintained on an isocaloric control diet or exposed to liquid diets containing 1.7% (PAE-L) or 3.4% (PAE-M) ethanol from gestational day (GD) 11 to GD19, followed by collection of mammary tissue from newborn female offspring for transcriptomic analysis (Fig. 1A). Unsupervised analysis of the transcriptomic profiles revealed broad differences in gene expression among control and ethanol-exposed mammary tissues. **The heatmap revealed distinct patterns of gene expression among the three groups, with the most prominent differences observed in the PAE-M group** (Fig. 1B). Consistent with this pattern, differential expression analysis identified both up- and downregulated genes following PAE, with substantially more transcriptional alterations in PAE-M than in PAE-L mammary tissue (Fig. 1C). Together, these findings demonstrate that prenatal ethanol exposure produces an exposure-level-dependent reprogramming of the newborn mammary transcriptome. We therefore next examined the individual differentially expressed genes and their patterns of regulation to identify molecular features underlying the PAE-associated transcriptional response.

**Figure 1.**
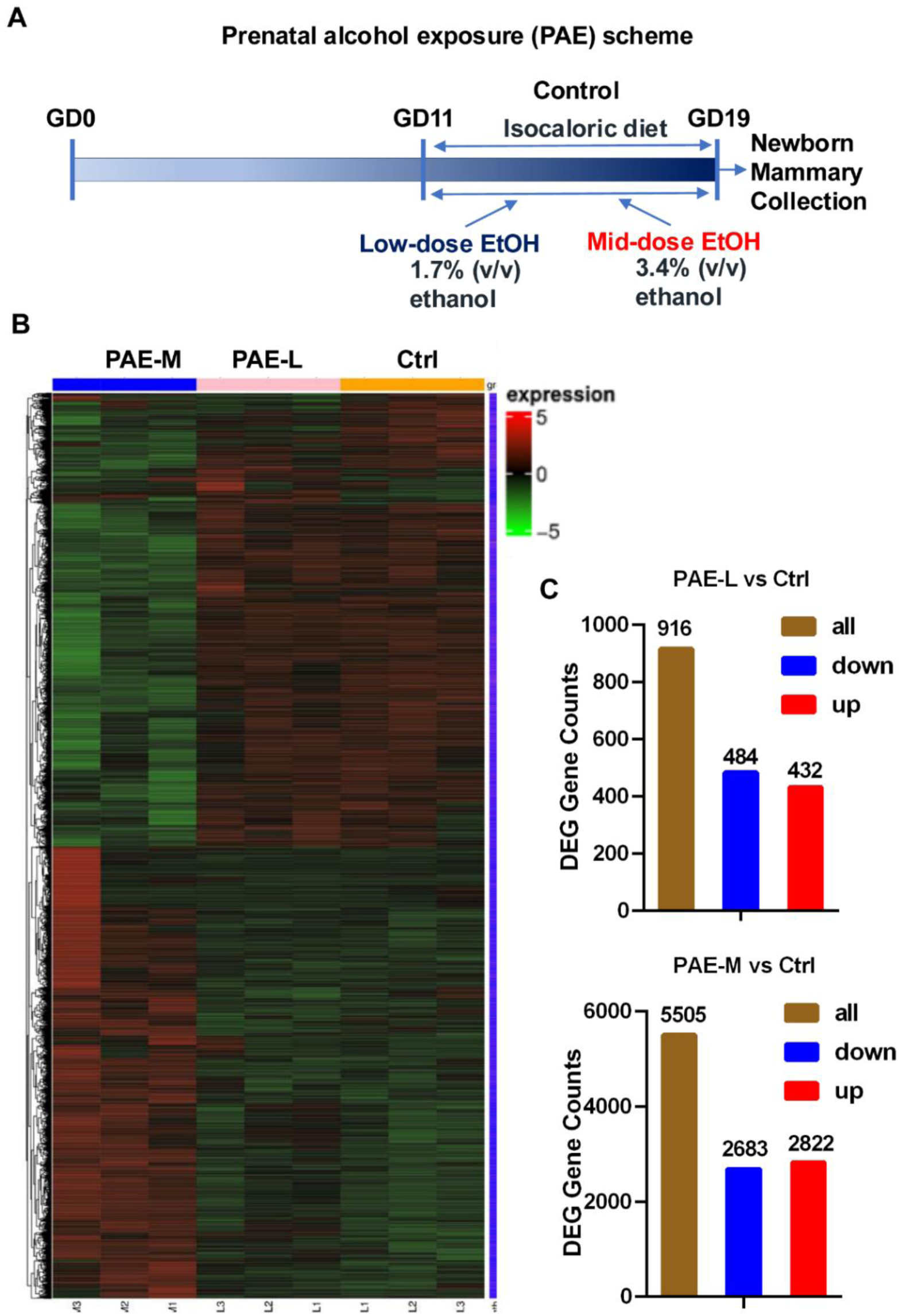
Experimental design and global transcriptomic profiles of newborn mammary tissues following prenatal ethanol exposure. (A) Experimental design. Pregnant mice received an isocaloric control diet or ethanol-containing liquid diets (PAE-L, 1.7% v/v; PAE-M, 3.4% v/v) from gestational day (GD) 11 to GD19. Mammary tissues were collected from newborn female offspring for RNA-seq analysis. (B) Heatmap showing global gene-expression profiles in control, PAE-L, and PAE-M mammary tissues. Expression values are displayed as row-standardized values according to the indicated color scale. (C) Numbers of total, upregulated, and downregulated differentially expressed genes (DEGs) for PAE-L versus control and PAE-M versus control comparisons using the indicated differential-expression criteria.

### 2. Prenatal ethanol exposure produces exposure-level-dependent patterns of differential gene expression

To further characterize the transcriptional response to prenatal ethanol exposure, differential gene expression was examined separately in PAE-L and PAE-M mammary tissues relative to controls. Volcano plot analysis revealed both increased and decreased gene expression following PAE, with marked differences in the magnitude and extent of the response between the two exposure conditions (Fig. 2A,B). PAE-L produced a relatively limited transcriptional response, whereas PAE-M resulted in substantially broader changes involving large numbers of both up- and downregulated genes.

**Figure 2.**
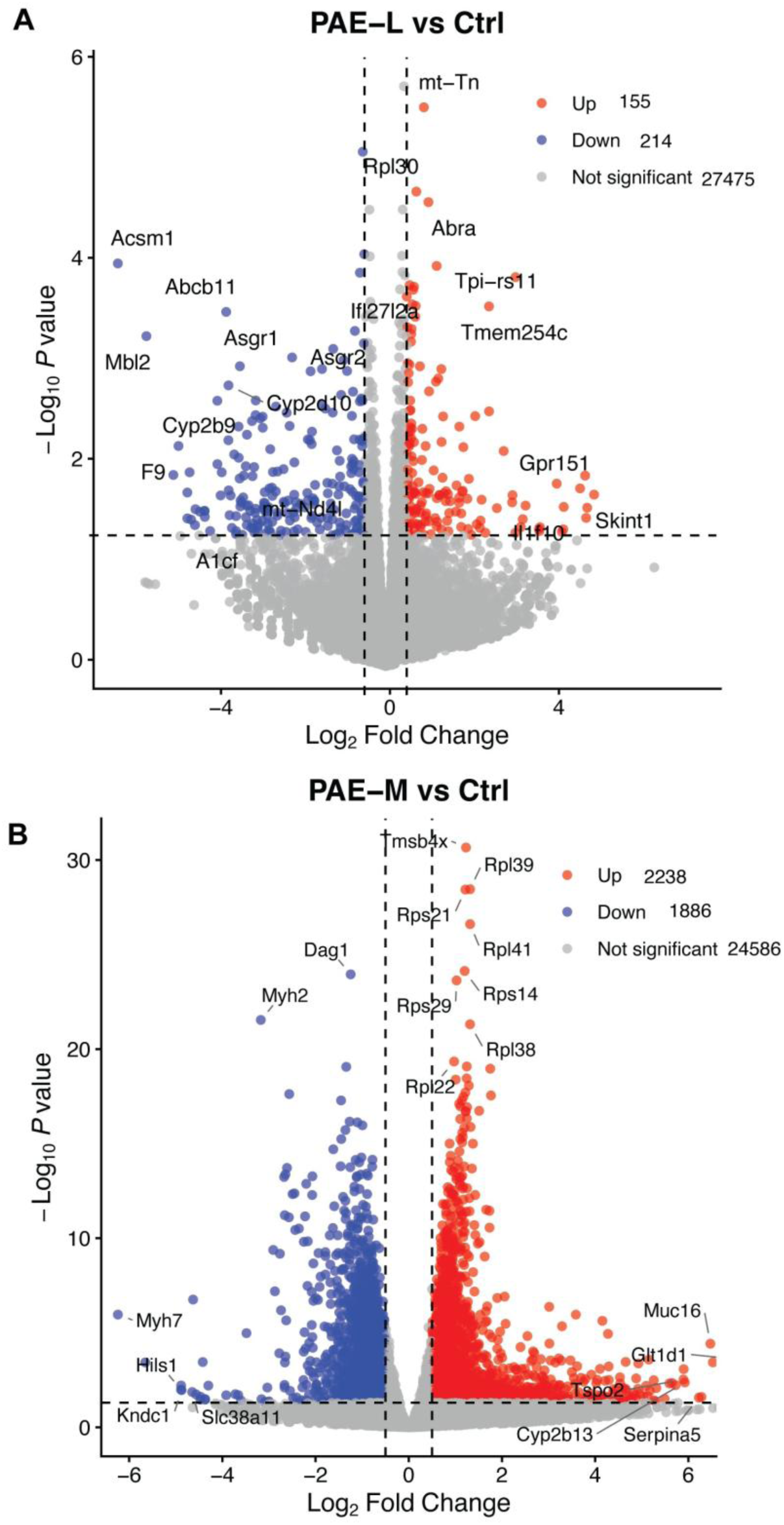
Differential gene-expression profiles in newborn mammary tissues following prenatal ethanol exposure. Volcano plots showing differentially expressed genes in (A) PAE-L versus control and (B) PAE-M versus control mammary tissues. Each point represents an individual gene plotted according to log2 fold change (log2FC) and statistical significance. Upregulated and downregulated genes are indicated by the colors shown, and selected highly altered genes are labeled. Numbers of upregulated and downregulated genes meeting the indicated differential-expression criteria are shown in each panel. The top 20 upregulated and downregulated genes for PAE-L and PAE-M relative to control are presented in Tables 1 and 2, respectively.

**Table 1.**
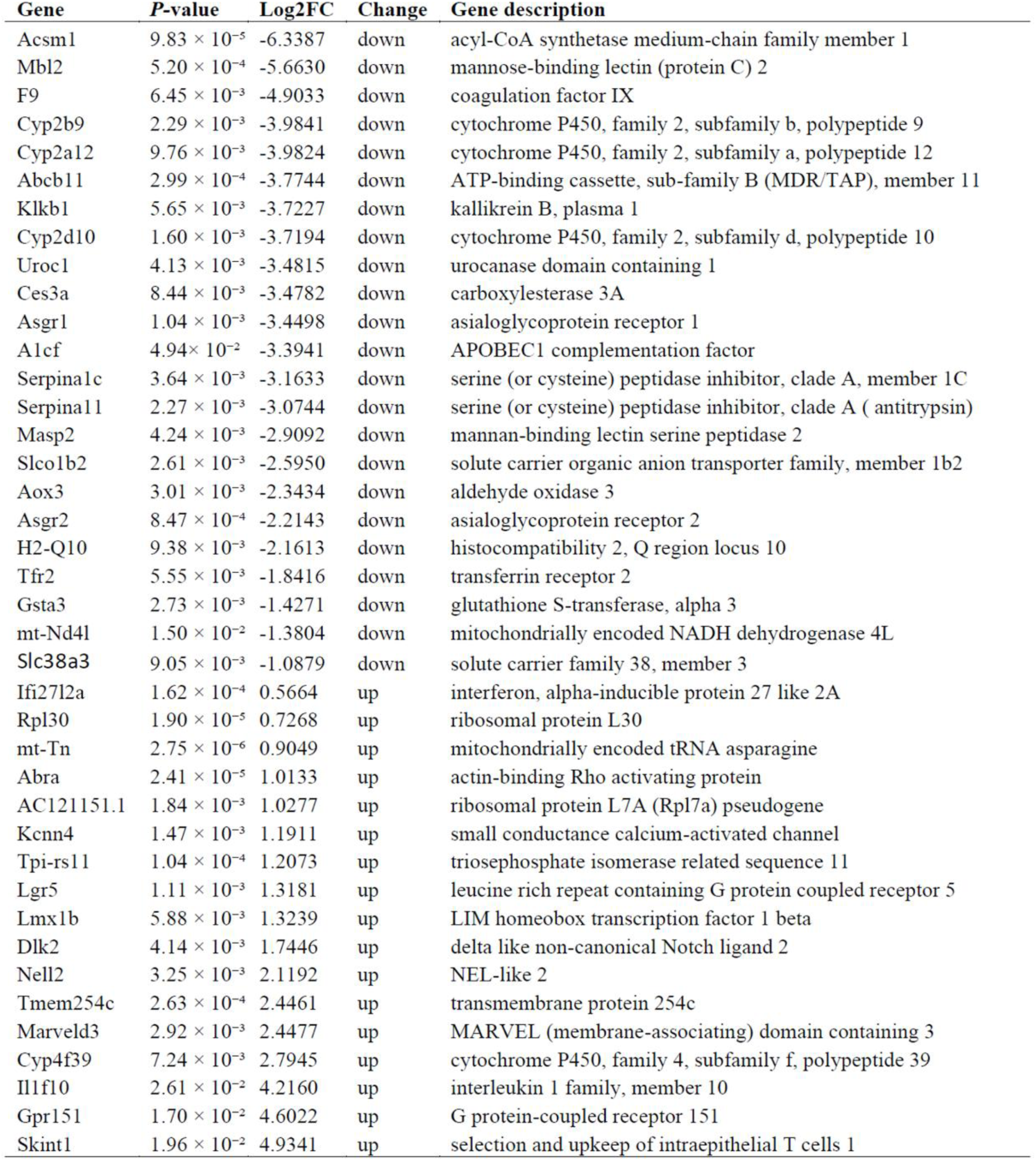
Top Differentially Regulated Genes in the PAE-L Group.

**Table 2.**
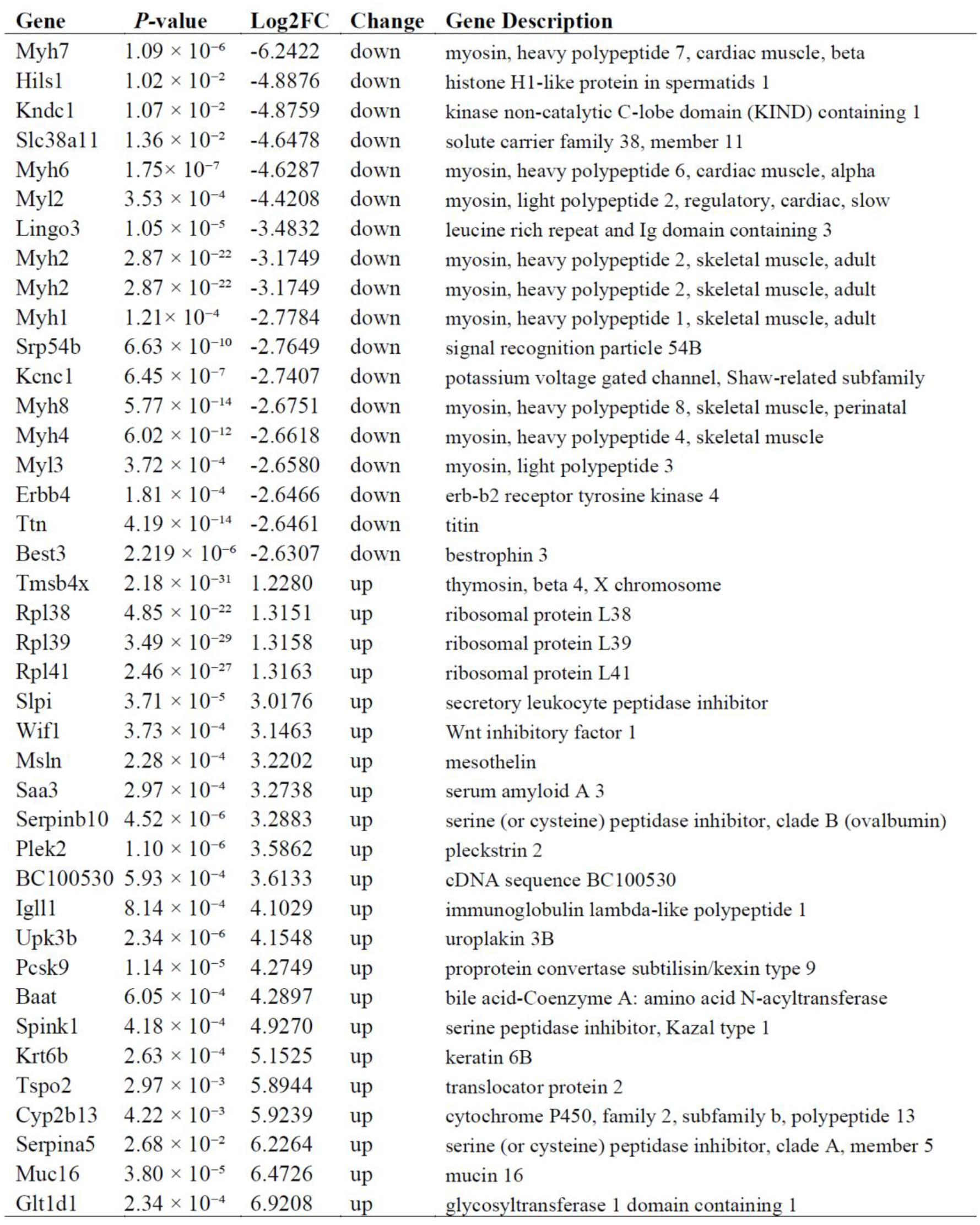
Top Differentially Regulated Genes in the PAE-M Group.

Examination of the most highly altered genes further demonstrated distinct transcriptional profiles between the two exposure groups (Tables 1 and 2). In PAE-L mammary tissues, prominent changes included decreased expression of genes involved in metabolic and transport functions, together with increased expression of genes with diverse regulatory functions, including Lgr5, Dlk2, Lmx1b, and Il1f10. In contrast, PAE-M produced a broader pattern that included marked alterations in structural, metabolic, signaling, and regulatory genes. Notably, Erbb4, a receptor tyrosine kinase with established roles in mammary gland development, was among the prominently downregulated genes in the PAE-M group. Other highly altered transcripts included Wif1, Pleckstrin 2 (Plek2), Muc16, and several ribosomal protein genes, further illustrating the diversity of the transcriptional response.

Together, these results demonstrate that prenatal ethanol exposure produces distinct exposure-level-dependent patterns of gene regulation in the newborn mammary gland, with PAE-M eliciting substantially more extensive transcriptional reprogramming than PAE-L. These findings prompted further pathway-level analyses to identify the biological processes associated with the PAE-induced transcriptional changes.

### 3. Prenatal ethanol exposure remodels insulin and metabolic signaling programs in the newborn mammary gland

To identify biological pathways associated with PAE-induced transcriptional changes, KEGG pathway enrichment analysis was performed separately for PAE-L and PAE-M mammary tissues relative to controls. PAE-L was associated with alterations in several metabolic and signaling pathways, including insulin signaling, insulin resistance, glucagon signaling, FoxO signaling, PPAR signaling, and steroid and retinol metabolism (Fig. 3A). PAE-M produced a broader and distinct pattern of pathway alterations involving metabolic, growth-factor, and cellular signaling processes (Fig. 3B). Despite the differences between the two exposure groups, **insulin signaling was identified in both analyses**, suggesting that this pathway represents a common target of prenatal ethanol exposure.

**Figure 3.**
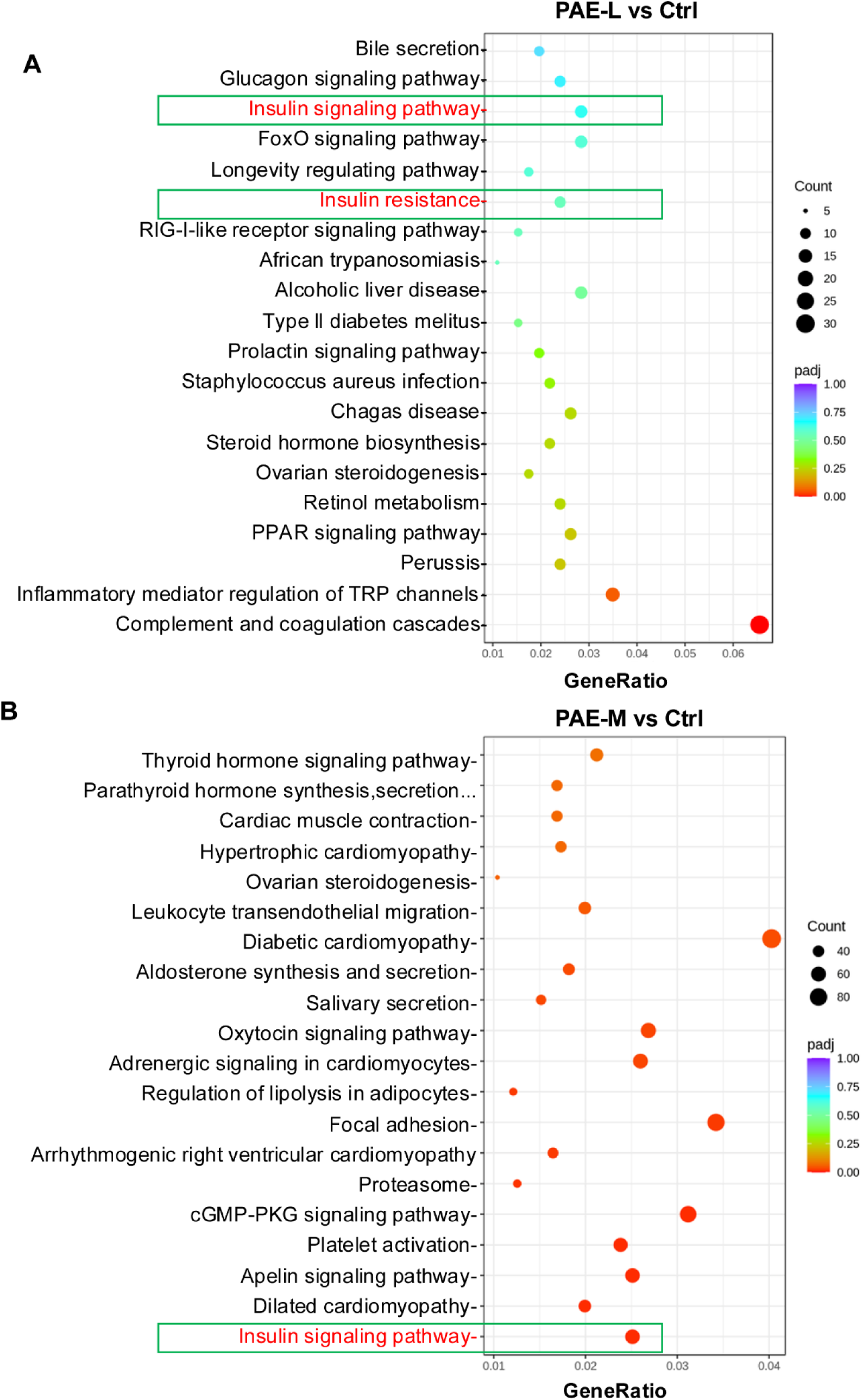
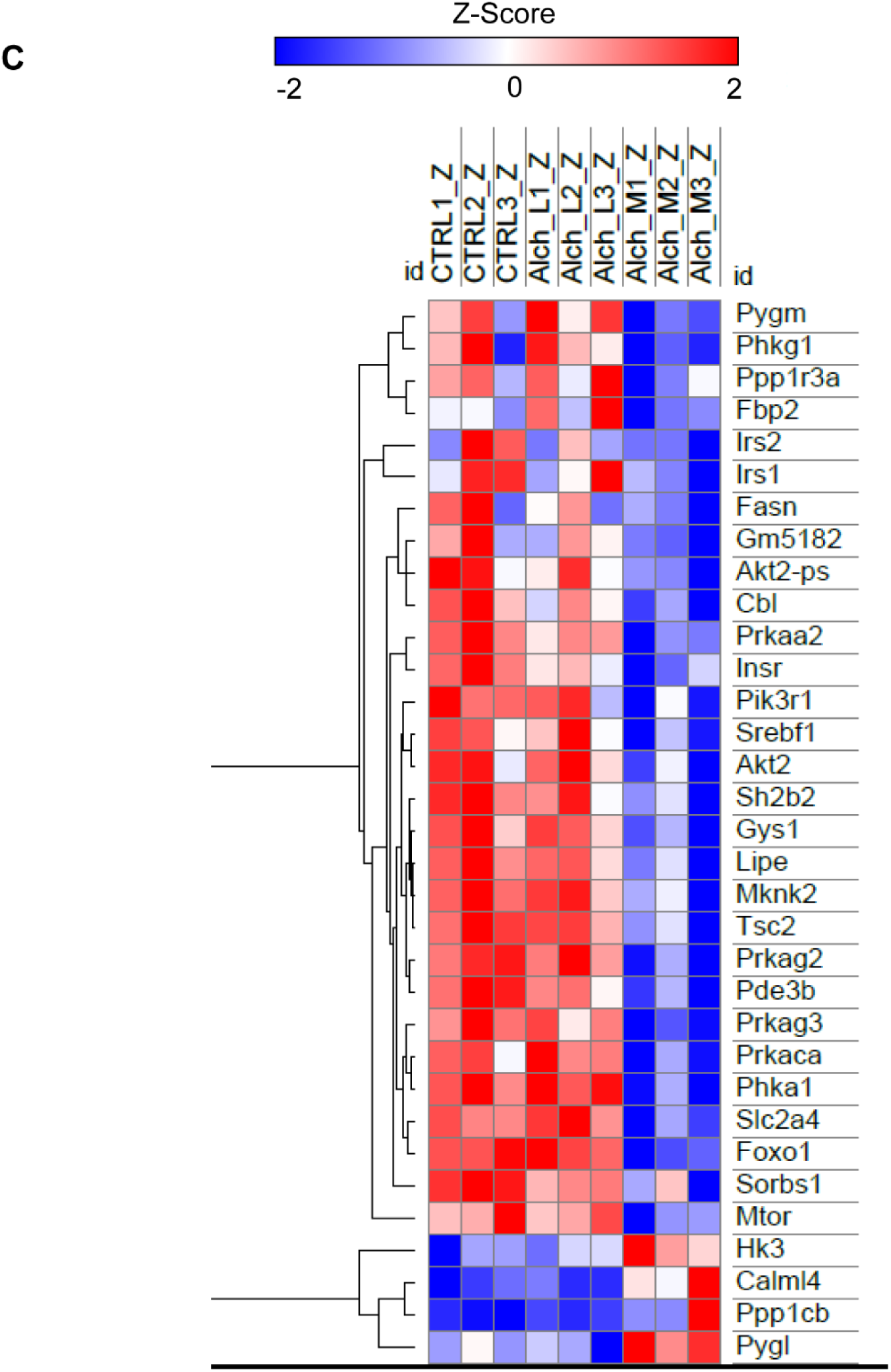
KEGG pathway analysis and insulin-signaling gene-expression profiles following prenatal ethanol exposure. **(A, B)** KEGG pathway enrichment analysis of differentially expressed genes in PAE-L versus control (A) and PAE-M versus control (B) newborn mammary tissues. Dot size represents the number of genes associated with each pathway, and color indicates the adjusted P value (padj). Insulin signaling is highlighted. **(C)** Heatmap of differentially expressed genes contributing to the KEGG insulin signaling pathway (mmu04910). Of the 58 genes contributing to pathway enrichment, genes meeting adjusted P < 0.05 and |log2 fold change| ≥ 0.5 in the PAE-M versus control comparison are displayed. Expression values were standardized by gene (row Z-score), genes were hierarchically clustered.

Because the PAE-M group exhibited the more extensive transcriptional response, we further examined genes contributing to the KEGG insulin signaling pathway (mmu04910). Among the 58 genes contributing to pathway enrichment, genes meeting adjusted *P* < 0.05 and |log2 fold change| ≥ 0.5 in PAE-M versus control were analyzed by hierarchical clustering (Fig. 3C). The resulting heatmap revealed coordinated alterations across multiple components of the insulin-signaling network. Most of the selected genes showed decreased expression in PAE-M mammary tissue, whereas a smaller subset was increased, indicating broad remodeling rather than uniform suppression of the pathway.

Together, these findings identify insulin/metabolic signaling as a prominent component of the transcriptional response to prenatal ethanol exposure in the newborn mammary gland. We therefore next examined selected components of this signaling network at the mRNA and protein levels.

### 4. Prenatal ethanol exposure alters IRS1-associated growth and metabolic signaling in the newborn mammary gland

To determine whether the transcriptomic changes in insulin/metabolic pathways were accompanied by alterations in related molecular signaling, selected components of the insulin/IGF and growth-factor signaling networks were examined in newborn mammary tissues. Expression of **IGFBP3** and **IGFBP5** mRNA was increased most prominently in PAE-L mammary tissues, with more modest changes observed following PAE-M exposure (Fig. 4A). These findings further indicated that the molecular response to PAE differed between the two exposure levels rather than following a simple linear exposure-response pattern.

**Figure 4.**
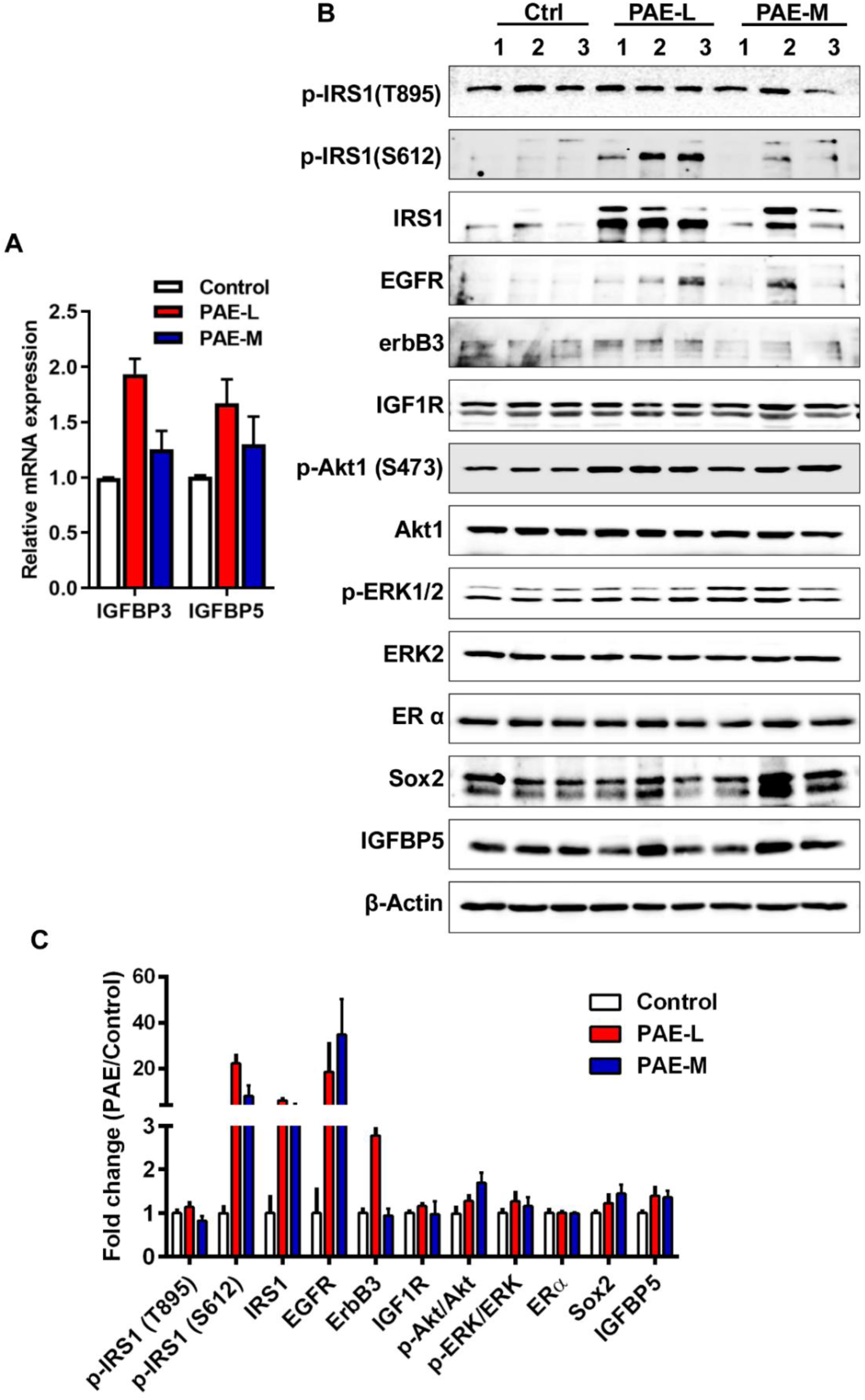
Effects of prenatal ethanol exposure on insulin/IGF- and growth-factor-associated signaling in newborn mammary tissues. **(A)** Relative mRNA expression of IGFBP3 and IGFBP5 in control, PAE-L, and PAE-M newborn mammary tissues determined by quantitative RT-PCR. **(B)** Representative Western blots of selected insulin/IGF- and growth-factor-associated signaling proteins, including p-IRS1(T895), p-IRS1(S612), IRS1, EGFR, ErbB3, IGF1R, p-Akt1(S473), Akt1, p-ERK1/2, ERK2, ERα, Sox2, and IGFBP5. **(C)** Densitometric analysis of Western blot data. Phosphorylated proteins were normalized to their corresponding total proteins, and total protein levels were normalized to β-Actin, as appropriate.

Protein analysis revealed particularly prominent alterations in IRS1-associated signaling (Fig. 4B,C). Total IRS1 abundance was markedly increased in PAE-L mammary tissue and was accompanied by a pronounced increase in IRS1 Ser612 phosphorylation, whereas phosphorylation at Thr895 showed comparatively little change. IRS1 abundance and phosphorylation in PAE-M tissues displayed a different pattern, further demonstrating exposure-level-dependent regulation of this signaling node.

Additional components of growth-factor signaling exhibited more variable responses. IGF1R abundance remained relatively stable, whereas EGFR and ErbB3 showed more pronounced but variable changes following PAE. In comparison, alterations in AKT and ERK phosphorylation were modest. Changes were also observed in proteins associated with mammary growth and differentiation, including ERα, Sox2, and IGFBP5 (Fig. 4B,C). Together, these findings demonstrate that prenatal ethanol exposure is associated with remodeling of IRS1-centered growth and metabolic signaling in the newborn mammary gland, with distinct molecular responses at the two exposure levels.

### 5. Prenatal ethanol exposure enhances proliferative and cell-cycle programming in the newborn mammary gland

To further define the biological processes associated with PAE-induced transcriptional reprogramming, Reactome pathway enrichment analysis was performed. In PAE-M mammary tissues, prominent enrichment was observed for pathways associated with RNA metabolism and translation as well as cell-cycle progression, including G1/S transition, S phase, DNA synthesis and replication, mitotic metaphase and anaphase, and separation of sister chromatids (Fig. 5A). In comparison, PAE-L exhibited a distinct enrichment profile dominated by RNA metabolism and multiple processes associated with translation initiation and protein synthesis (Fig. 5B).

**Figure 5.**
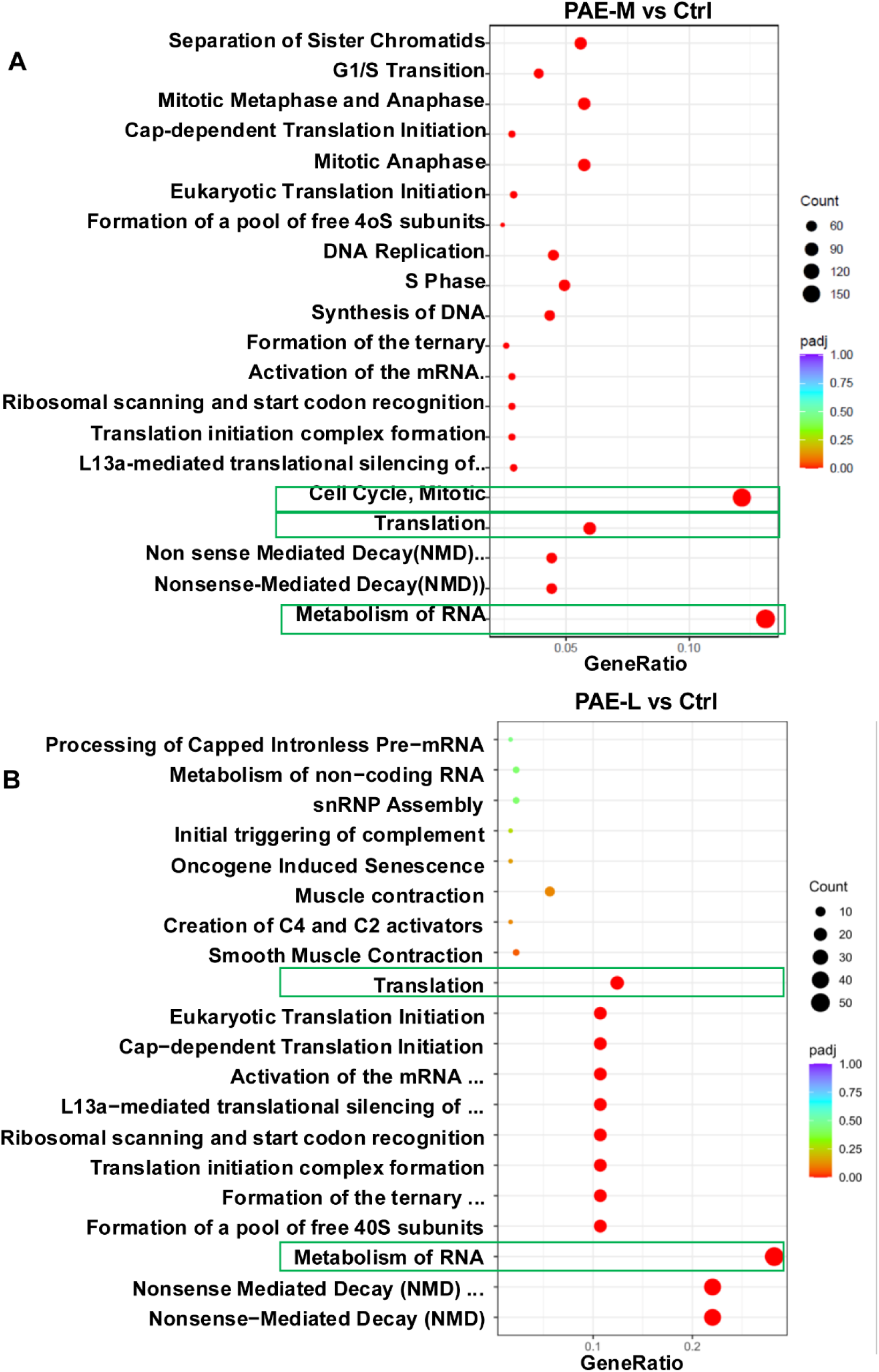
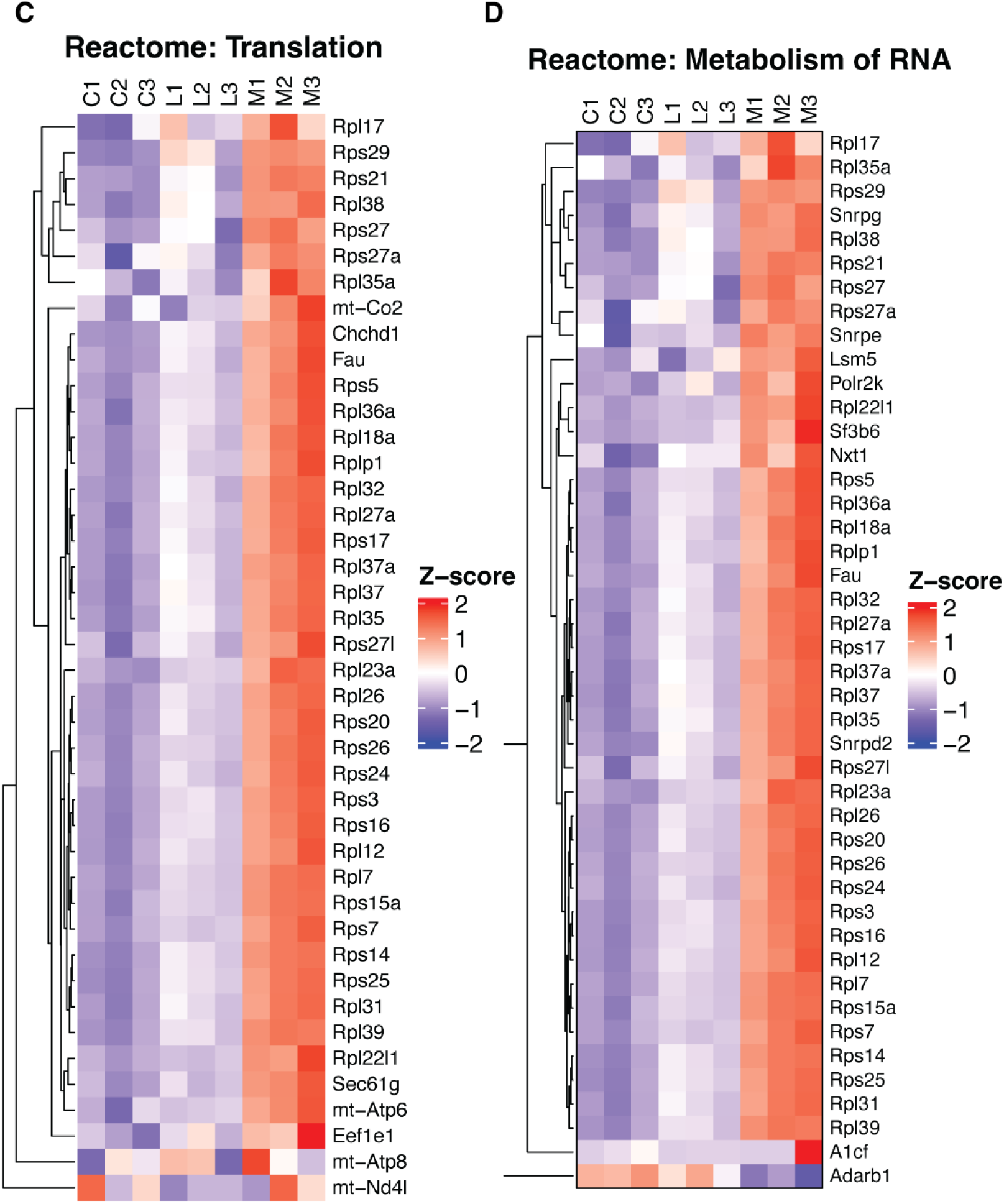
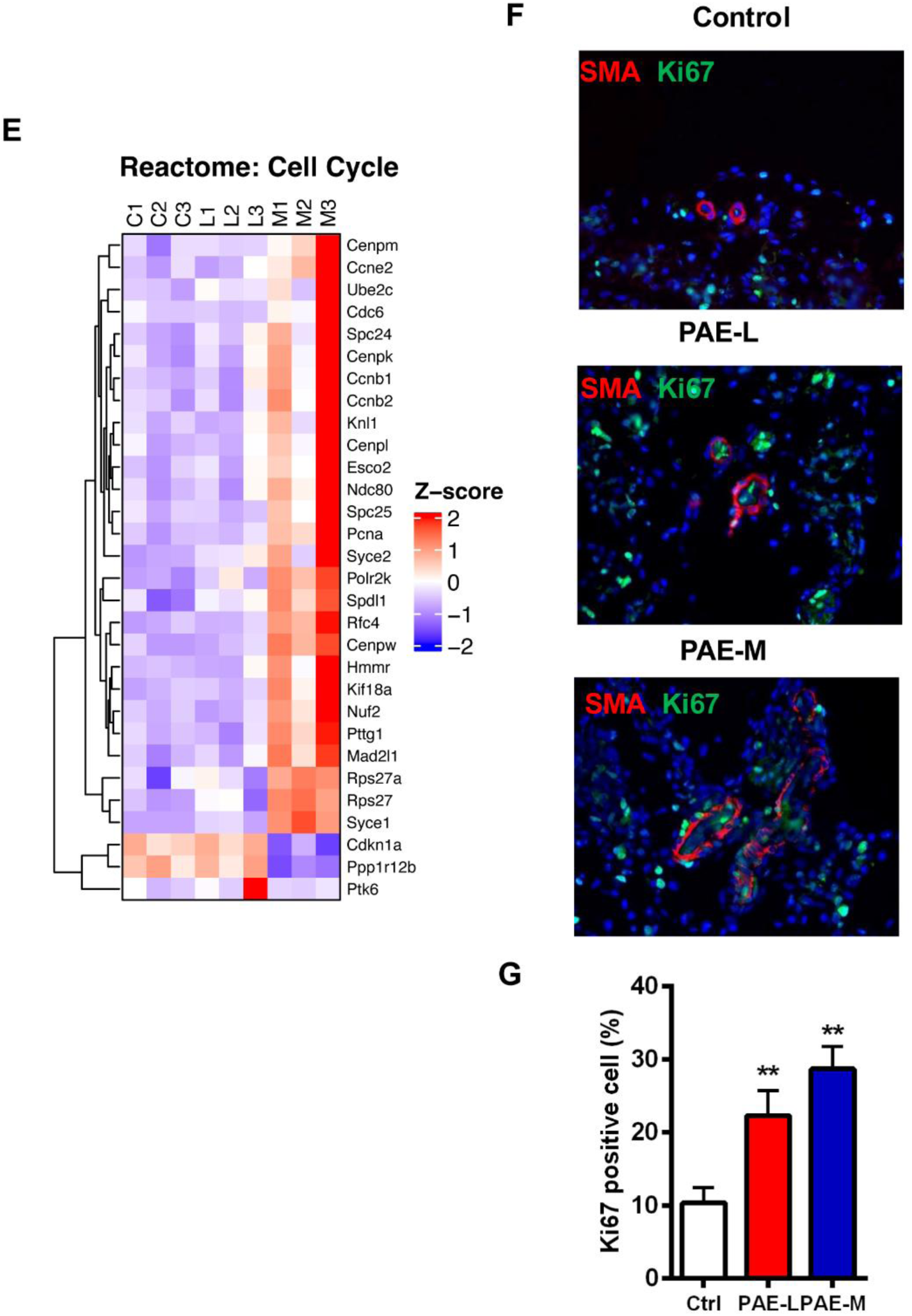
Prenatal ethanol exposure alters RNA/translation programs and enhances cell-cycle and proliferative activity in the newborn mammary gland. **(A, B)** Reactome pathway enrichment analysis of differentially expressed genes in PAE-M versus control (A) and PAE-L versus control (B) newborn mammary tissues. Dot size represents the number of genes associated with each pathway, and color indicates the adjusted P value (padj). Enriched pathways include processes associated with RNA metabolism, translation, DNA replication, and cell-cycle progression. **(C–E)** Focused heatmaps of differentially expressed genes associated with RNA metabolism, translation, and mitosis/cell-cycle programs. Expression values were standardized by gene (row Z-score), and genes were hierarchically clustered. Sample order was fixed by experimental group (control, PAE-L, and PAE-M). **(F)** Representative immunofluorescence images of newborn mammary tissues from control, PAE-L, and PAE-M offspring stained for α-smooth muscle actin (SMA, red), Ki67 (green), and nuclei (DAPI, blue). **(G)** Quantification of Ki67-positive cells within mammary tissue.

To examine these transcriptional programs in greater detail, focused heatmaps were generated for differentially expressed genes associated with RNA metabolism, translation, mitosis, and cell-cycle regulation (Fig. 5C–E). These analyses revealed coordinated upregulation of genes within these functional programs rather than changes restricted to individual transcripts. RNA metabolism and translation-associated genes were increased following PAE, while genes involved in mitosis and cell-cycle progression showed a particularly prominent increase in PAE-M mammary tissues. The cell-cycle gene set included multiple genes involved in DNA replication, chromosome segregation, and mitotic progression, consistent with the pathway-level enrichment observed in the Reactome analysis.

To determine whether the transcriptional signatures associated with cell proliferation were accompanied by a tissue-level phenotype, newborn mammary tissues were examined by immunofluorescence for α-smooth muscle actin (SMA) and Ki67 (Fig. 5F). Ki67-positive cells were more abundant in mammary tissues from both PAE-L and PAE-M offspring compared with controls. Quantification confirmed a significant increase in the percentage of Ki67-positive cells following both exposure conditions, with the highest level observed in PAE-M mammary tissues (Fig. 5G).

Together, these findings demonstrate that prenatal ethanol exposure alters RNA-processing and translational programs and is associated with enhanced cell-cycle and proliferative activity already evident in the newborn mammary gland. The concordance between the transcriptional signatures and increased Ki67-positive cells provides evidence that the PAE-associated molecular reprogramming is accompanied by an early proliferative phenotype.

## Discussion

The present study demonstrates that prenatal alcohol exposure is associated with substantial molecular reprogramming of the mammary gland that is already evident at birth. Transcriptomic profiling of newborn mammary tissues revealed exposure-level-dependent changes in gene expression, with the more extensive response observed following the moderate exposure condition. Pathway analyses identified two particularly prominent features: remodeling of insulin/metabolic signaling and alterations in programs associated with RNA metabolism, translation, DNA replication, mitosis, and cell-cycle progression. These transcriptional changes were accompanied by altered IRS1-associated signaling and increased Ki67-positive cells in newborn mammary tissue. Together, these findings indicate that the fetal mammary gland responds directly to the prenatal exposure environment and that PAE establishes an altered molecular and proliferative state before extensive postnatal mammary development has occurred.

These findings extend previous observations from our laboratory and others showing persistent effects of PAE on mammary development later in life [15–17, 21]. In the MMTV-ErbB2 model, prenatal ethanol exposure altered mammary morphogenesis and growth-regulatory signaling and was associated with increased mammary tumor susceptibility [15]. Our more recent studies in a non-tumorigenic background further demonstrated persistent changes in postnatal mammary morphogenesis, epithelial proliferation, mammary epithelial populations, stem/progenitor-associated activities, and growth-regulatory signaling [21]. The current study moves the developmental window backward and demonstrates that substantial molecular changes are already detectable in the newborn mammary gland. These early changes provide a potential developmental foundation upon which subsequent postnatal endocrine, stromal, metabolic, and environmental influences may act. Thus, the prenatal and postnatal effects of PAE should not be viewed as mutually exclusive processes; rather, molecular changes initiated during fetal development may be compounded with additional developmental and/or external factors as the mammary gland undergoes postnatal growth and differentiation.

A prominent finding from the transcriptomic analysis was the involvement of insulin and related metabolic signaling. Insulin signaling was identified in the pathway analyses of both exposure conditions, and focused analysis in PAE-M tissues demonstrated coordinated changes in multiple components of this network, including genes associated with receptor/IRS signaling, PI3K–AKT/mTOR regulation, and downstream metabolic control. The pattern was not consistent with simple uniform activation or suppression of the pathway, but instead suggested broader remodeling of insulin/metabolic regulation. This distinction is important because insulin/IGF signaling has multiple functions during mammary development, influencing epithelial growth, survival, differentiation, and interactions with other growth-factor pathways [26–28]. Alteration of this network during fetal mammary development could therefore affect the developmental state of the tissue without necessarily producing a single directional signaling response.

The biochemical findings further support this interpretation. Changes in IGFBP3 and IGFBP5 expression were accompanied by pronounced alterations in IRS1 abundance and phosphorylation, particularly in the PAE-L group. Total IRS1 and IRS1 Ser612 phosphorylation increased markedly, whereas other components, including IGF1R and downstream AKT and ERK signaling, exhibited more modest or variable responses. The stronger IRS1 protein response in PAE-L, despite broader transcriptional changes in PAE-M, also emphasizes that the biological response to PAE is not simply proportional to exposure level. Differences between transcript abundance, protein abundance, phosphorylation, feedback regulation, and turnover may all contribute to the observed patterns. Accordingly, these data are most consistent with exposure-dependent remodeling of the insulin/IGF–IRS signaling network, rather than straightforward activation or inhibition of a single pathway.

The individual gene-expression data also provide potentially informative links to mammary developmental regulation. For example, Erbb4 was among the prominently altered genes. ERBB4 is a member of the ErbB receptor family with established roles in mammary epithelial differentiation and development, making its alteration notable in the context of the newborn mammary gland [29]. Lgr5, another gene altered following PAE, has been associated with mammary epithelial stem/progenitor populations and developmental lineage states [30]. These observations do not establish that PAE directly alters a specific mammary stem-cell population, particularly because the present RNA-seq analysis was performed on whole dissected newborn mammary tissue. Nevertheless, they raise the possibility that some of the early transcriptional changes identified here may involve developmental programs that influence epithelial lineage composition or stem/progenitor activity. This possibility is particularly relevant to our previous postnatal observations and warrants direct investigation using cell-type-resolved approaches.

A second major feature of the transcriptomic response involved RNA metabolism, translation, and proliferative programs. Reactome analysis identified RNA metabolism and translation-related processes following both exposure conditions, while PAE-M additionally showed prominent enrichment of pathways involving G1/S transition, DNA replication, S phase, chromosome segregation, and mitosis [16]. Focused gene-expression analyses further demonstrated coordinated increases across these functional programs. Altered RNA processing and translational regulation may be especially relevant during a rapidly developing tissue state, in which changes in protein synthesis capacity can accompany shifts in cellular growth and differentiation. Importantly, the enrichment of cell-cycle and mitotic programs was supported by an independent tissue-level observation: quantitative analysis demonstrated significantly increased Ki67-positive cells in mammary tissues from both PAE-L and PAE-M offspring [25]. Thus, the transcriptional signatures associated with proliferation correspond to a measurable proliferative phenotype already present at birth.

The insulin/metabolic and proliferative programs were emphasized here because they represented particularly prominent and biologically relevant features of the PAE response, but they constitute only part of the information contained within the transcriptomic dataset. The global expression and pathway analyses identified numerous additional signaling, metabolic, developmental, structural, and regulatory processes that may provide leads for subsequent investigation. Further analysis of these pathways, together with integration of the present dataset with later developmental stages, may help distinguish transient responses from molecular changes that persist during postnatal mammary development. In addition, approaches that preserve spatial and cellular information, including spatial transcriptomic profiling and cell-type-resolved analyses, could determine whether specific PAE-responsive programs originate primarily from mammary epithelial cells, surrounding mesenchymal/stromal populations, or interactions among these compartments.

This consideration is particularly relevant to the newborn mammary gland. At this developmental stage, the mammary epithelial structure is small and embedded within a complex surrounding tissue environment, and bulk RNA-seq necessarily captures contributions from multiple cellular compartments. Some prominent transcriptional changes, including genes associated with muscle or contractile functions, may therefore reflect changes in surrounding mesenchymal or other adjacent cell populations as well as changes intrinsic to the mammary epithelium. Rather than separating these possibilities with the present dataset, the findings highlight the need to consider the developing mammary gland as an integrated tissue environment. Future spatial and cell-specific studies should help resolve the cellular origins of the transcriptional responses identified here.

Overall, our findings establish that the effects of prenatal ethanol exposure on mammary development are detectable at the molecular and cellular levels by birth. PAE produces broad, exposure-level-dependent transcriptional reprogramming involving insulin/metabolic signaling, RNA and translational regulation, and cell-cycle and proliferative programs, accompanied by altered IRS1-associated signaling and increased mammary cell proliferation. These early changes do not necessarily determine the subsequent developmental trajectory by themselves; rather, they define an altered developmental state that may interact with the extensive endocrine, cellular, and environmental changes occurring during postnatal mammary development. The newborn mammary gland therefore provides an important window for understanding how prenatal exposure initiates developmental changes that may contribute to persistent mammary phenotypes later in life.

## Data Availability Statement

All data is contained within the manuscript.

## Author Contributions

ZM: Data curation and analysis, writing-review & editing; YL: Data curation and analysis, writing-review & editing; AP: Data curation and analysis, writing-review & editing; XY: Conceptualization, funding acquisition, investigation, project administration, writing – drafting, editing & review. All authors have read and agreed to the published version of the manuscript.

## Funding

This work was supported in part by a R16 grant from the National Institute of General Medical Sciences (1R16GM145545) to XY, a U54 grant from the National Institute on Alcohol Abuse and Alcoholism (U54 AA019765), and a RCMI U54 grant from the National Institute on Minority Health and Health Disparities (U54 MD012392).

## Acknowledgments

The authors extend their appreciation to the funding agency and support from colleagues in this department.

## Conflict of Interest

The authors declare no conflicts of interest.

## Notes

### Competing Interest Statement

The authors have declared no competing interest.

